# Docking for molecules that bind in a symmetric stack with SymDOCK

**DOI:** 10.1101/2023.10.27.564400

**Authors:** Matthew S. Smith, Ian S. Knight, Rian C. Kormos, Joseph G. Pepe, Peter Kunach, Marc I. Diamond, Sarah H. Shahmoradian, John J. Irwin, William F. DeGrado, Brian K. Shoichet

## Abstract

Discovering ligands for amyloid fibrils, such as those formed by the tau protein, is an area of much current interest. In recent structures, ligands bind in stacks in the tau fibrils to reflect the rotational and translational symmetry of the fibril itself; in these structures the ligands make few interactions with the protein but interact extensively with each other. To exploit this symmetry and stacking, we developed SymDOCK, a method to dock molecules that follow the protein’s symmetry. For each prospective ligand pose, we apply the symmetry operation of the fibril to generate a self-interacting and fibril-interacting stack, checking that doing so will not cause a clash between the original molecule and its image. Absent a clash, we retain that pose and add the ligand-ligand van der Waals energy to the ligand’s docking score (here using DOCK3.8). We can check these geometries and energies using an implementation of ANI, a neural network-based quantum-mechanical evaluation of the ligand stacking energies. In retrospective calculations, symmetry docking can reproduce the poses of three tau PET tracers whose structures have been determined. More convincingly, in a *prospective* study SymDOCK predicted the structure of the PET tracer MK-6240 bound in a symmetrical stack to AD PHF tau before that structure was determined; the docked pose was used to determine how MK-6240 fit the cryo-EM density. In proof-of-concept studies, SymDOCK enriched known ligands over property-matched decoys in retrospective screens without sacrificing docking speed, and can address large library screens that seek new symmetrical stackers. Future applications of this approach will be considered.

## Introduction

An open problem in ligand discovery is understanding and exploiting the ability of high-affinity ligands to bind to protein amyloids. Tau (tubulin associated unit), an intrinsically disordered protein that can wrap around microtubules, is one such target for diagnostic tools.^1–3^ The accumulation of tau fibrils into toxic neurofibrillary tangles (NFTs) is characteristic of Alzheimer’s disease (AD), chronic traumatic encephalopathy (CTE), and other neurodegenerative tauopathies.^4–10^ In cryogenic electron microscopy (cryo-EM) studies, tau adopts different polymorphs characteristic of different diseases: the paired helical filament (PHF) and the straight filament (SF) for AD and Type I or Type II for CTE.^11–17^ Recently, cryo-EM structures of AD tau and CTE tau with ligands bound have been determined. A striking feature of these structures is that the ligands bind in stacks to the tau filaments to reflect the rotational and translational symmetry of the fibril. Epigallocatechin gallate (EGCG), a tau disaggregator found in green tea, binds at the inter-protofilament cleft of AD PHF tau with a stoichiometry of one ligand molecule per protein monomer.^18^ The positron emission tomography (PET) radiotracers GTP-1 and MK-6240 bind within the “C” shape of each protofilament with the same stoichiometry, as does flortaucipir (Tauvid) to CTE Type I tau.^19–22^

In each of these examples, the ligands within the stacks make substantial van der Waals and π-π interactions. Intriguingly, for the PET tracers, these ligand-ligand interactions seem more dominant than their direct contacts with the fibrils. For example, GTP-1 forms only a single hydrogen bond with lysine 353 on AD PHF tau.^19^ Flortaucipir’s pyridine nitrogen in its benzo-pyrrolo-pyridine moiety makes a hydrogen bond to aspartate 358 in CTE Type I tau; however, apart from this, the structure primarily extends into the solvent.^21^ EGCG establishes more hydrophilic contacts with the inter-protofilament cleft of AD PHF tau, including hydrogen bonds to asparagine 327, glutamate 338, and lysine 340, and polar contact with histidine 329. Yet, its interactions mainly involve other ligands in the stack and the surrounding solvent.^18^ Despite these minimal fibril contacts, GTP-1 exhibits an 11 nM affinity to tau, while flortaucipir binds to brain-derived tau in the low nM to high pM concentration range.^23–30^

To use this new mode of binding to find other ligands that might bind to different tau polymorphs and to other protein fibrils, we developed the symmetric extension to DOCK3.8^31–33^, “SymDOCK.” This method uses the same pose sampling and scoring as in DOCK3.8 but it only retains those poses that can form a symmetrical stack in the protein. We then evaluate the ligand-ligand van der Waals energy as a crude measure of favorable self-interaction and add that to the DOCK score. Given the imposed symmetry, we assume that the interactions with the protein are copied from ligand monomer to monomer. We found that in both retrospective and, more convincingly, in prospective prediction, SymDOCK succeeded in recapitulating and in predicting the poses of different tau ligands. In a test-of-concept screen of 22 million molecules, SymDOCK was not much slower than base DOCK3.8. This suggests that the extra cost of the new symmetry operations and of calculating inter-ligand van der Waals energies is compensated by the lower sampling imposed by the symmetry constraints. To further test SymDOCK’s generated geometries, we computed their energies using ANI, a neural network force field trained on density functional theory (DFT) energy data.^34,35^ We found that their ANI energies and exact geometries differed only slightly from ANI-optimized stacks. We consider further generalizations of this approach.

## Methods

### Symmetric Pose Sampling

The SymDOCK extension uses the same ligand building and initial pose generation as base DOCK3.8, allowing a user to dock the built section of the ZINC22 database without any modifications.^31,36^ To keep SymDOCK from decreasing the speed of docking each molecule, we do not generate or attempt to dock a stack of molecules. Rather, we generate poses (conformations and orientations) for each ligand as always in DOCK, but only keep those that can self-stack.^31,37,38^ Because of the symmetry of the system, we only need to evaluate the ligand-protein interaction (checks for clashes and energetic favorability) for a single site; the ligand-protein interactions at all other sites will be identical.

Once we generate a potential pose, we apply the symmetry operation of the fibril and check the ligand-ligand interactions (**Fig. 1a-b**). The symmetry operation of the fibril is the 4-dimensional affine transformation matrix that acts on a vector in 3 dimensions as the application of a rotation followed by a translation. For the tau fibril, this transformation would be a 4.7 Å translation down the long axis of the fibril along with a ∼-1° rotation in the plane perpendicular to that axis.^11,18^ We describe a procedure for generating this matrix in general in the discussion of **Supplemental Fig. 1**. Once we have a rotated/translated pose of the molecule, we check for a van der Waals clash with the initial pose, defined as any atom from the initial pose being within 2 Å of any atom in the transformed pose. We choose this conservative definition of what constitutes a clash based on the experimental structure of GTP-1 bound to AD PHF tau (**Supplemental Fig. 2**).^19^ For each pose that passes the symmetry check, we add the van der Waals interaction of the original molecule and the transformed version to the total DOCK score function, with the same AMBER 4.0 united-atom force field parameters as the ligand-protein van der Waals energy.^31,39–41^

**Figure 1.**
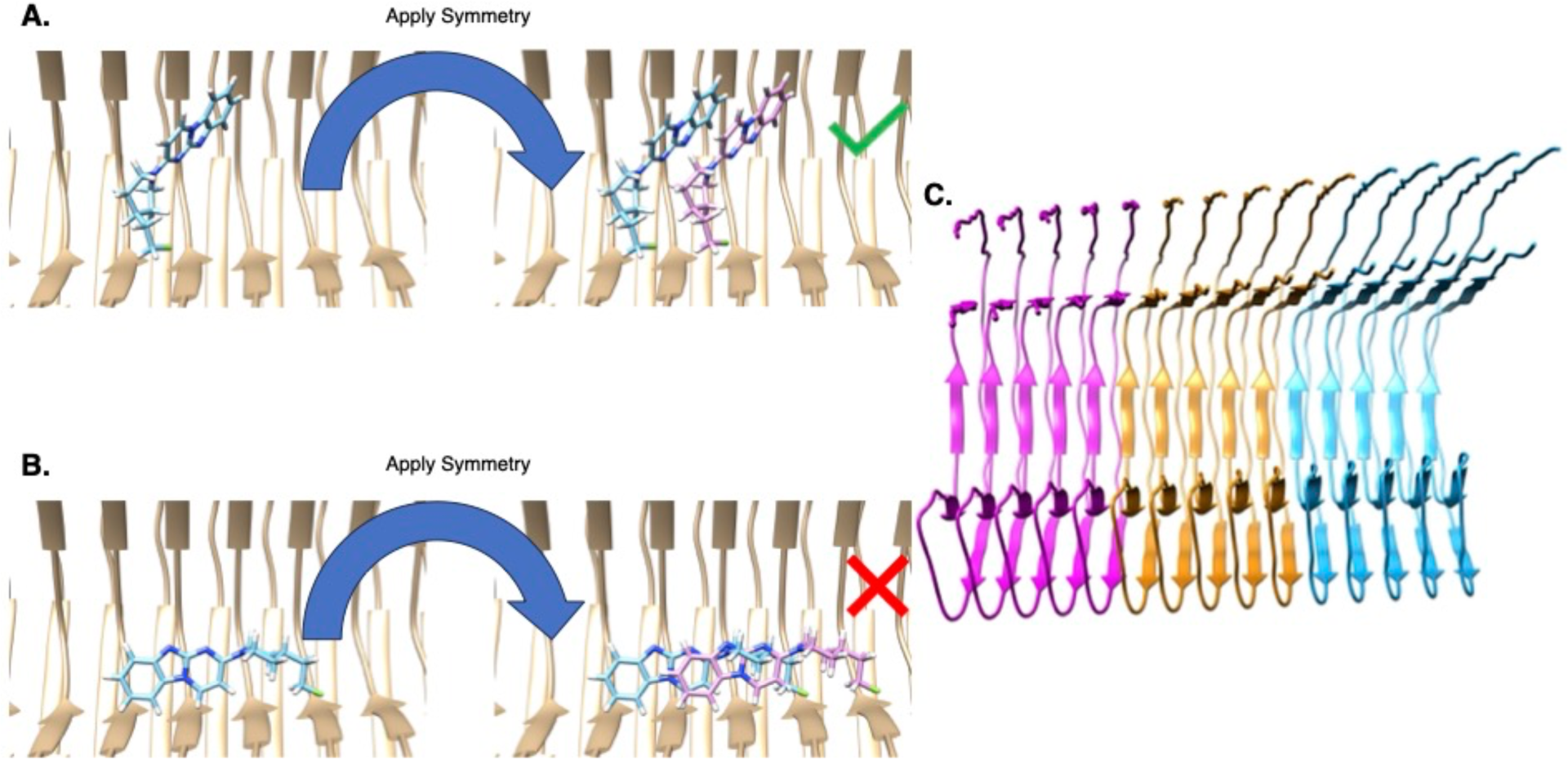
A demonstration of SymDOCK by docking GTP-1 into the structure of it bound to AD PHF tau (PDB ID: 8FUG). **a-b.** SymDOCK generates poses for molecules as DOCK 3.8 normally does and then applies a symmetry check: if the molecule (blue) and its translated/rotated copy (purple) have no atoms within 2 Å of each other, the pose passes the filter (**a.**, with the dummy example of the experimental pose). If the molecule and its copy have atoms within this distance, the pose fails the filter (**b.**, showing the top pose of docking without the symmetry requirement). SymDOCK only evaluates ligand-ligand van der Waals energies of poses that pass the filter, and only does so for the first pairs’ energies, since ligand-protein interactions will be the same at every site by symmetry (ligand-protein interaction terms are evaluated as in DOCK3.8 for the first ligand pose in the stack). **c.** To avoid edge effects from modeling a small fraction of a fibril that should be nanometers in length, we artificially extend the input protein structure using the symmetry operation. This cryo-EM structure only contains 5 monomers of tau in each protofilament, but by applying the affine transformation and its inverse on the monomer atom positions we can generate new atom positions for a fibril of 15 monomers (bronze for original structure, blue for applying symmetry operation, and magenta for applying the inverse symmetry operation).

### Changes to the DOCK Score Grids

In addition to the explicit ligand-ligand interactions, we slightly change how we calculate the grids for the ligand-protein interactions. Since the protein fibril is hundreds of nanometers long, its interactions with any small molecule ligand should be periodic.^11^ We simulate this by artificially extending the fibril structure to more monomers than what is in the deposited structure (**Fig. 1c**). For example, the cryo-EM structure of GTP-1 bound to AD PHF tau (PDB ID: 8FUG; we used an earlier version of the structure) only contains 5 monomers of tau in each protofilament. By applying the affine transformation and its inverse on the monomer atom positions, we can generate new atom positions for a fibril of 15 monomers. We then feed the longer fibril into the Pydock3 procedure^42^ for generating the van der Waals, electrostatic, and desolvation grids and see that these grids do not have edge effects in the region of interest to docking (**Supplemental Fig. 3**).

We also change the electrostatic and desolvation grids to account for the changed dielectric environment of a stack of organic ligand molecules (**Supplemental Fig. 4**). Part of the normal DOCK3.8 optimization procedure includes adding a low dielectric layer that extends the boundary of protein atoms into the bulk solvent.^31,39,42^ This increases the magnitude of electrostatic interactions, attempting to balance the non-polar terms in docking, which often dominate. In SymDOCK, we have two sources of large-magnitude van der Waals energies: ligand-protein interactions and ligand-ligand interactions. Extending the low dielectric boundary for both electrostatic and for desolvation grids to the positions of the heavy atoms in a stack of ligands in the experimental structure helps to overcome this bias toward non-polar terms and mimics the ligand binding’s effect on the dielectric boundary (**Supplemental Fig. 5**). Such a ligand perturbation to the dielectric boundary becomes even more important with a stack of ligands bound to the fibril.

### ANI-Based Stack Filter

The energetic favorability of the molecular poses returned by SymDOCK may be screened using the ANI-2x molecular force field, as implemented in TorchANI.^34,35^ ANI-2x is a neural network trained to predict the ground state energies of small organic molecules, as calculated by density functional theory at the ωB97X/6-31G* level. We reasoned that the ANI-2x force field provided the best compromise between throughput and accuracy for the goal of determining the per-monomer ground state energies of the stacks of small molecules generated by SymDOCK.

We first estimate the SymDOCK-predicted geometry’s per-monomer energy, then use that as a starting geometry for a Monte Carlo optimization via the Metropolis-Hastings algorithm.^43^ We sample each candidate move by taking translations from a multivariate normal distribution and rotations from an isotopic normal distribution on SO(3) (the group or rotations in 3D).^44^ For each candidate move, we generate the two nearest symmetry mates of the molecule, evaluate the ANI energy, and determine the per-monomer energy (**Supplemental Methods**). After 1,000 Monte Carlo steps, we compare the minimum energy configuration achieved to SymDOCK’s output both in terms of ANI-generated energy difference and RMSD. We do not alter the conformation of the molecule from what SymDOCK produces.

## Results

### Retrospective Pose Reproduction

Reproducing experimental poses of known binders was a first metric we used to evaluate SymDOCK. Typically, reproducing the ligand pose by docking it back into its native complex is considered necessary in the field, but here there is the complication of regenerating the full ligand stack. We began with the PET tracer GTP-1, docking it back into its complex with the AD PHF tau fibril. As is typical, we used pseudo atoms (“spheres”) nucleated around the known ligand coordinates to define the sampling, and calculated van der Waals, Poisson-Boltzmann electrostatic, and desolvation energy potential grids to evaluate ligand protein complementarity.^31,39,45^ SymDOCK sampled GTP-1 in 1507 orientations in the fibril site, and multiple conformations within each orientation.^46^ Each pose was checked for the ability to generate a symmetry mate that does not clash with itself using the fibril symmetry operator (**Fig. 1a**). If a pose passed this check, then SymDOCK calculated the van der Waals, electrostatic, desolvation and, new to SymDOCK, ligand-ligand van der Waals energies. We then ranked allowed poses by total DOCK score. When we compared the highest-scoring docked pose of GTP-1 to its experimental geometry, the root-mean-squared difference (RMSD) was 1.165 Å, with the three-ring system of the ligand almost exactly superposed and the differences coming from the flexible tail (**Fig. 2a, Supplemental Fig. 6**). Similarly, docking EGCG back into its complex in the inter-protofilament cleft of AD PHF tau led to an RMSD of 2.91 Å to the experimental structure for the best-fitting pose (**Fig. 2b**). This pose was the second-best according to DOCK score (ignoring poses that differed only in placement of phenolic hydroxyl hydrogens), and the highest-scoring one had an RMSD of 6.83 Å from the experimental structure (**Supplemental Fig. 7**). Despite the RMSD of the best fitting, 2.91 Å pose, SymDOCK recapitulated most of the key polar interactions observed in the experimental structure, including hydrogen bonds to Asn327 and His329 on one side of the cleft and Glu338 and Lys340 on the other. Just as importantly, all the predicted poses captured the ligand-ligand aromatic stacking. Finally, docking flortaucipir back into its complex with CTE Type I tau led to an RMSD of 0.477 Å for the best docking pose, 7.78 Å for the top scoring one, and RMSD values between of 7.49Å and 7.74 Å for the poses scoring between those two (**Fig. 2c, Supplemental Fig. 8**). When we compared GTP-1’s, EGCG’s and flortaucipir’s SymDOCK poses to their ANI-optimized geometries, the ligand-ligand energies improved but the structures changed only between 0.6 and 1.1 Å in RMSD (**Supplemental Table 1).** Crucially, in all three of the docked complexes the ligand-ligand packing and quadrupole stacking closely resembled those in the experimental structures.

**Figure 2.**
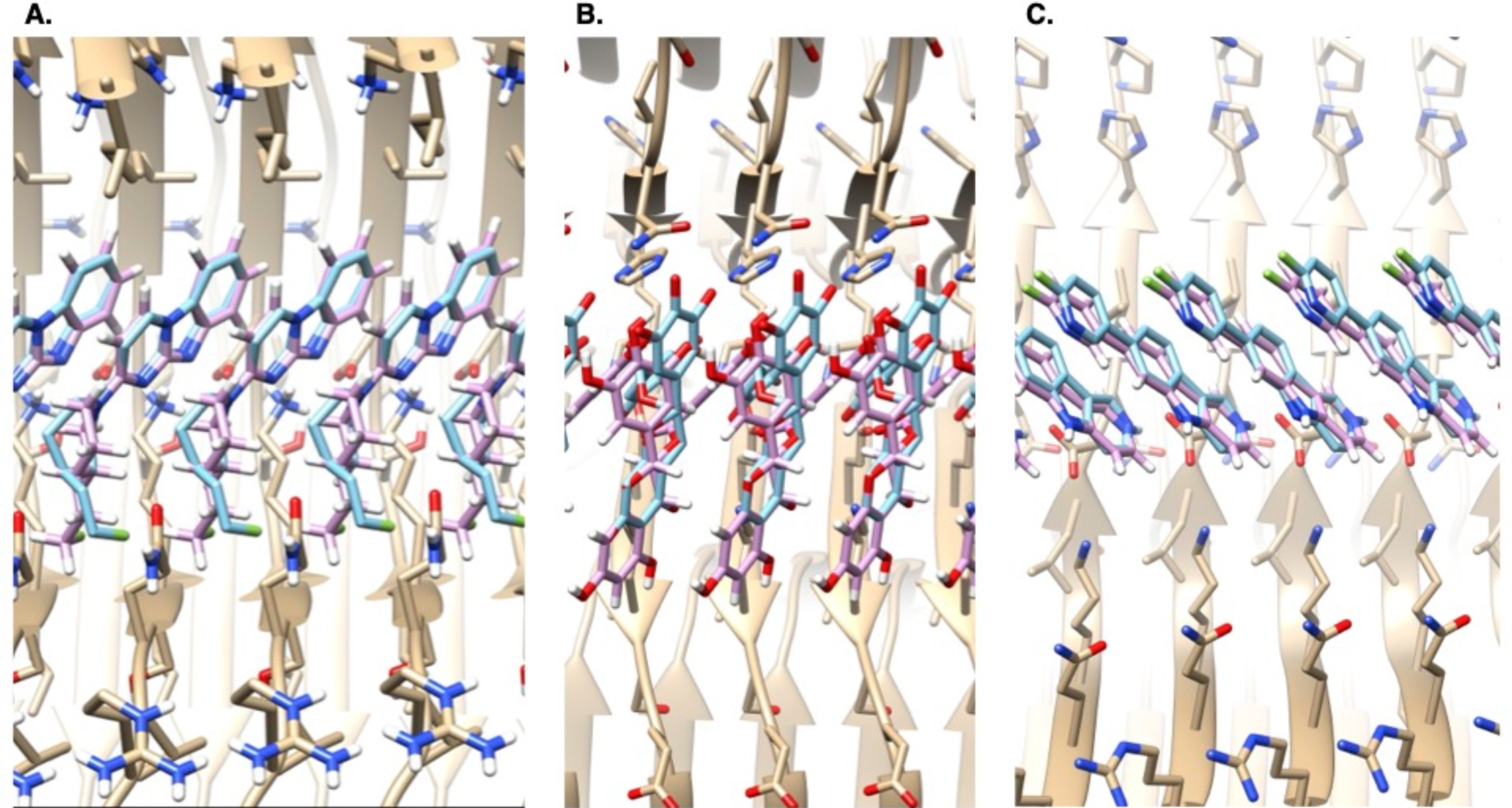
Retrospective pose reproduction, with the protein in beige, the cryo-EM pose in blue, and the SymDOCK-predicted pose in purple. **a.** GTP-1 docked into in the PET-tracer-site of AD PHF tau, RMSD to the experimental structure 1.2 Å. **b.** EGCG docked into the inter-protofilament cleft of AD PHF tau, RMSD to the experimental structure 2.9 Å. **c.** Flortaucipir docked into CTE Type I tau, RMSD to the experimental 0.48 Å.

### Retrospective Enrichment of Ligands Over Decoys

We were curious if the symmetry docking approach could identify plausible fibril ligands from among a large molecular library. We began by asking whether SymDOCK could prioritize known ligands versus a much larger set of property-matched decoys. Docking against AD PHF tau, we docked the PET ligands GTP-1, MK-6240, and a set of 14 MK-6240 analogs, drawn from a larger group of 94 to maximize diversity (**Supplemental Table 2**).^20,47^ We generated 50 property matched decoys for each ligand, for 800 total.^48,49^ Docking the 16 ligands and 800 decoys against the GTP-1-bound structure of AD PHF tau resulted in an adjusted logAUC enrichment of 32.7 and an enrichment factor at 1% (EF_1%_) of 6.4 (**Supplemental Fig. 9**).^39,50^ The better ranking of the known ligands versus the decoys reflects their better ability to form stacks that recapitulate the fibril symmetry and can interact with it. For instance, ZINC000000035542 is a 1.2.2-bicyclo that can dock to the receptor grids using the non-symmetric default DOCK3.8 procedure, but its 3-dimensional ring structure prevents it from docking symmetrically (structure in **Supplemental Fig. 10**). Even though this decoy has properties matched to the known ligands, the symmetry check prevents it from scoring and improves enrichment.

### Docking 22 Million Molecules from ZINC22

Some of the decoys from the enrichment calculation scored well and looked plausible, making good stacking interactions with themselves and packing in the fibril to mimic its symmetry. We wondered if there might be many molecules in a general library that might be suitable to form symmetrical stacks in a fibril. To test how symmetry docking would work in a large library screen, we docked 22 million molecules from the ZINC22 database^36^ against the GTP-1-bound form of AD PHF tau. We selected molecules that were similar in gross properties to GTP-1: neutral, with heavy atom count between 21 and 23, and clogP values between 3.30 and 3.50.^51,52^ The 22 million molecules docked in 17,800 core hours (less than a day on a typical 1000-core cluster), 2.90 seconds per molecule per core which is two to three times slower than typical for standard docking with DOCK3.8.^32^ When passed through the ANI-based filter, the configurations of the stacked ligands typically changed only modestly (**Supplemental Fig. 11**), and the energy changes were similar to what we had seen with the PET ligands (**Supplementary Table 1**). Visually, many of these top 5000 molecules stacked in plausible ways, typically making one or two polar interactions with the fibril, akin to what is observed in the experimental ligand-fibril structures. For instance, ZINCnD000001AHba appears to make a single hydrogen bond to Lys353 from its amide carbonyl (**Fig. 3a**). The molecule stacks with a symmetry reflecting that of the overall fibril, and the 4.7 Å translation ensures that the putative quadrupole-quadrupole interactions that the benzothiophene and quinoline rings make are weak,^53–55^ but with water exclusion still likely contributing to a hydrophobic effect of stacking. Meanwhile, ZINCmD000004I6kF and ZINCnF000004BXys are not posed to form explicit hydrogen bonds with the fibril, instead making van der Waals contacts with it and, perhaps more importantly, making extensive ligand-ligand stacking interactions (**Fig 3b-c**). Self-stacking also seems to dominate in the pose of the amide-linked phenyl-benzimidazole ZINClD000002Tbny, which again makes no formal hydrogen bond to the fibril (**Fig. 3d**). When docked to the same site, the known PET ligands GTP-1 and MK-6240 would have placed among the top 5 to 7% best-scoring molecules in this screen of 22 million (the ranking of the other 14 known ligands used in the retrospective study are in **Supplemental Table 3**). The poses of the molecules taken from ZINC22 lack experimental testing and should not carry too much weight, but they do support the plausibility of large library screens using a symmetry-based ligand stacking approach.

**Figure 3.**
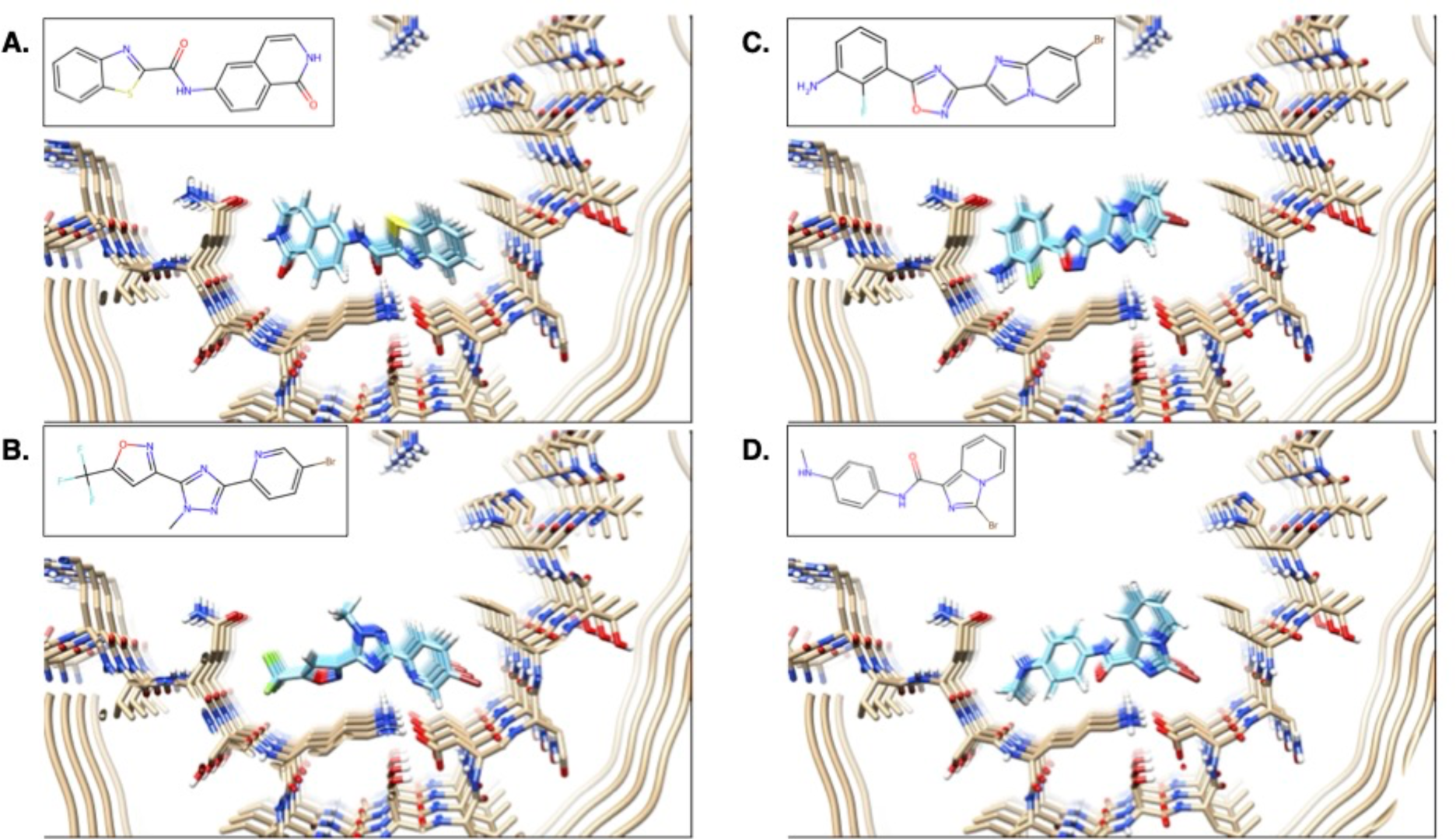
Stacking and polar interactions to the AD PHF tau (beige) among docked poses of four representative high-ranking molecules from a 22-million-molecule library screen (blue). **a.** The benzothiophene-quinoline ZINCnD000001Ahba. **b and c.** The triaryls ZINCmD000004I6kF and ZINCnF000004Bxys. **d.** The amide-linked phenyl-benzimidazole ZINClD000002Tbny.

### Prospective Pose Prediction

Encouraged by the retrospective results and the ability to generate plausible geometries from large library screens, we tried the method for genuine prospective pose prediction. Aware that the structure of the PET ligand MK-6240 was being determined against AD PHF tau fibrils, we docked that molecule against the structure of AD PHF tau in its GTP-1 complex using SymDOCK. When we compared the predicted docked poses to the subsequently-determined cryo-EM electron density, we found that the 9^th^-best scoring pose came close to fitting the electron density (**Fig. 4a-b)** and captured the symmetry and the overall placement of the ligand stack. The one discrepancy was that the ligand stack extended out of the cryo-EM density in its pyrrolo-pyridine ring and did not completely fill it on the other side of the molecule, indicating a simple rigid-body translation of the SymDOCK pose could better fit the density. Indeed, comparing the final refined position for the ligand stack to that predicted by the docking, the RMSD of 2.13 Å can be mostly attributed to a rotation about the axis of the fibril (**Fig. 4c-d**). Indeed, the structure was close enough that the docked pose was used as the input for refinement of the final experimental structure of the MK-6240/AD PHF tau complex.^56^ We note that although this was the 9th-best scoring structure, the best pose by DOCK score ha an RMSD of 3.01 Å from the experimental structure, and the third-best scoring pose had an RMSD of 2.27 Å to the experimental structure.

**Figure 4.**
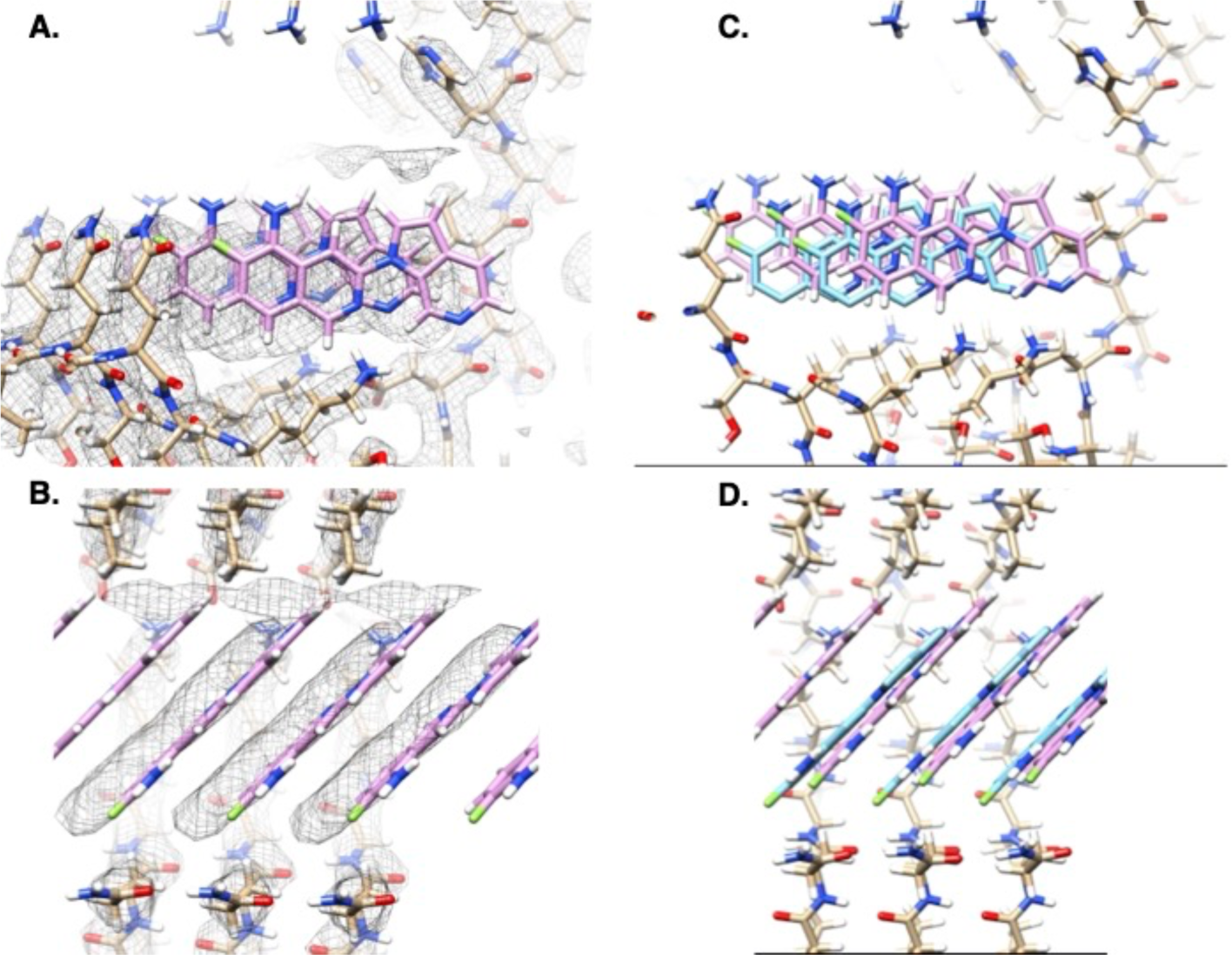
Prospective pose prediction for MK-6240 in AD PHF tau. **a-b.** The electron density of the MK-6240-tau structure (black mesh showing the 0.0117 level) with the modeled protein in beige and SymDOCK-predicted pose in purple. **a.** Looking down the axis of the fibril. **b.** Looking into the open trough of the fibril. **c-d.** The final refinement of the MK-6240 pose into the electron density (blue) using the SymDOCK-predicted pose as input.

## Discussion

An arresting feature of the recent tau fibril structures is the adoption of symmetrical stacks by their ligands (**Figs. 2 and 4**). Ligand-receptor interactions are few in these structures, with the most extensive contacts coming between the ligands themselves. For a receptor with multiple symmetrical sites, adoption of such ligand stacks may be the favorable mode. Considerations of entropy coming from the number of sites and cooperativity from the ligand-ligand interactions may explain how molecules with so few fibril interactions may nevertheless bind in the nM range, as many of the PET ligands do. A challenge for ligand discovery is exploiting the symmetrical fibril sites and ligand-ligand interactions, as most docking methods anticipate a 1:1 ligand-to-receptor stoichiometry and are designed to explore multiple ligand poses unencumbered by symmetry or stacking.

The SymDOCK approach adapts docking by imposing the symmetry of the fibril on any ligand geometry, insisting that there are no ligand-ligand van der Waals clashes. The method succeeds retrospectively in finding the experimental poses of four topologically unrelated PET ligands to tau (**Figs. 2 and 4**). While the symmetry calculation and the internal clash checks add operations to docking, these are compensated by the constraint imposed on pose selection by insisting on symmetry, and so the method loses little speed. A user can thus perform large library screens seeking new ligands. While these predictions remain to be tested experimentally, known fibril ligands are enriched both against property matched decoys (**Supplemental Fig. 9**) and in screens of 22 million diverse molecules (**Supplemental Table 3**). The new ligands’ poses seem plausible, forming stacks around planar aromatic and heteroaromatic systems while preserving the symmetry of the fibril (**Fig. 3**). In a genuine prospective prediction, the method correctly predicts the binding pose of MK-6240 to AD PHF tau (**Fig. 4**), and that docking pose was in fact used in the solution of the ligand-complex structure as determined by cryo-EM.^56^

Several limitations of SymDOCK merit airing. For speed of calculation, we have used a 2 Å cutoff to define ligand-ligand clashes, which misses those owing to large radii atoms like bromine or iodine. Only evaluating the favorability of a stacking geometry using the AMBER 4.0 united-atom force field’s van der Waals parameters ignores higher-order effects from π-π stacking and electrostatics.^40,41^ We have explored compensating for this with more detailed evaluation using the DFT-derived ANI,^34,35^ though this is slow enough to limit application to rescoring post-docking. We currently only use the symmetry operation (affine transformation) from the protein, but it is possible that other symmetries of a ligand stack could fit the pocket. For example, one could imagine ligands that bind in a 2:1 stoichiometry by forming stacks of molecules alternating in orientation with respect to the protein by 180° rotations. Finally, we have shown that prospective large library screens are mechanically possible using SymDOCK and produce plausible symmetrical stacks, but prospective hits and binding modes will demand experimental testing. For now, these predictions merely suggest that it should be possible to test this method for ligand discovery.

Notwithstanding these caveats, the key observations from this study should be clear. Imposing symmetry and excluding ligand-ligand clashes is an effective approach for docking ligands that form extensive, symmetric stacks against fibril proteins, and potentially other proteins with symmetrical sites. Because we used a relatively simple approach to this problem, the method is fast enough for use in large screens. Encouragingly, by insisting on symmetry and excluding clashes, the method seems to capture the key features of the symmetrical ligand stacks, despite ignoring higher-order energetic terms. We have implemented the symmetry docking approach in a program SymDOCK (freely available for academic research at https://dock.compbio.ucsf.edu/); we suspect that this approach may be readily adapted to most docking methods.^57–73^

## Author Contributions

The symmetry docking method was conceived of by MSS and BKS and implemented in DOCK3.8 by ISK under the supervision of JJI. MSS performed all docking calculations reported here. RCK developed the ANI-based stack filter under the supervision of WFD, and MSS used it to analyze the SymDOCK-derived poses. PK ran the experiments and computational analysis to provide the electron density and protein model for the structure of MK-6240 bound to AD PHF tau, under the supervision of MID and SHS, as described in an accompanying manuscript.^56^ All authors contributed to writing and editing the paper.

## Notes

The authors declare the following competing financial interest(s): B.K.S. serves on the SAB of Schrodinger, Vilya Therapeutics, and of Hyku Therapeutics, is a founder of Epiodyne, and with J.J.I of Deep Apple Therapeutics and BlueDolphin Leads LLC, and consults for Great Point Ventures and for Levator Therapeutics.

## Code Availability

SymDOCK is available without charge for academic research as part of the DOCK3.8 suite of programs at https://dock.compbio.ucsf.edu/.

