## Supplementary Materials for "Docking for molecules that bind in a symmetric stack with SymDOCK"

Contents:

Supplemental Methods

Supplemental Tables

- Supplemental Table 1
- Supplemental Table 2
- Supplemental Table 3

Supplemental Figures

- Supplemental Figure 1
- Supplemental Figure 2
- Supplemental Figure 3
- Supplemental Figure 4
- Supplemental Figure 5
- Supplemental Figure 6
- Supplemental Figure 7
- Supplemental Figure 8
- Supplemental Figure 9
- Supplemental Figure 10
- Supplemental Figure 11

**Supplemental Methods**

The energetic favorability of the molecular poses returned by SymDOCK may be screened using the ANI-2x molecular force field. ANI-2x is a neural network architecture trained to predict the ground state energies, as estimated by density functional theory with the ωB97X/6-31G* functional, of systems consisting of small organic molecules. We expected that the ANI-2x force field could provide the best compromise between throughput and accuracy for the goal of determining the per-monomer ground state energies of the stacks of small molecules generated by SymDOCK. The objective of the ANI-based screen was to determine whether each SymDOCK stack falls within user-specified thresholds of per-monomer energy and molecular pair RMSD of the optimal stack in the manifold of screw-symmetric molecular stacks with the same internal coordinates for each monomer. An approximate optimal stack is sampled via the Metropolis-Hastings algorithm on one molecule extracted from the SymDOCK stack and its two nearest symmetry mates. The affine transformation required to generate the stack is computed and independent translational and rotational MCMC moves are made to the stack during sampling. The translational moves are sampled from a Gaussian distribution, while the rotational moves are sampled from the isotropic Gaussian distribution on SO(3). Proposed moves are composed with the affine transformation matrix to generate a new matrix, which is then applied to the coordinates. Acceptance probabilities are determined using the Metropolis-Hastings criterion from the per-monomer ANI energies of the stack, with a temperature of 298 K. Per-monomer energies are computed from the many-body expansion of the stack energy up to trimer energies, assuming that only monomers, neighboring dimers, and trimers make contributions, as follows:

$$E_{per-monomer}=\frac{1}{3}\left( E_{trimer}-2E_{dimer}+3E_{Monomer} \right)+\left( E_{dimer}-2E_{monomer} \right)+E_{monomer}$$

$$=\frac{1}{3}\left( E_{trimer}+E_{dimer} \right)$$

While only molecular trimers are utilized when computing ANI energies, stacks of 10 consecutive molecules are generated for an additional long-range clash filter. If atoms from different molecules in these stacks come within a distance of 2Å, the proposed stack is rejected. Metropolis-Hastings MCMC sampling is run for a user-specified number of steps (1000 by default) and the minimal-energy stack is selected for calculation of the per-monomer energy difference with the SymDOCK stack and the pair-RMSD between adjacent pairs in the minimal-energy stack and the SymDOCK stack. SymDOCK stacks for which both the per-monomer energy difference and the pair-RMSD fall within the user-specified cutoffs advance through the screen.

**Supplementary Tables**

| **Molecule** | **pre_opt_energy (kcal/mol)** | **post_opt_energy (kcal/mol)** | **pre_post_opt_rmsd (Å)** |
| --- | --- | --- | --- |
| **GTP-1** | -7.110 | -9.158 | 0.574 |
| **EGCG** | -1.402 | -9.585 | 1.081 |
| **Flortaucipir** | -6.310 | -12.563 | 0.752 |

**Supplemental Table 1.** Comparing best the docked pose for GTP-1, EGCG, and flortaucipir against their ANI-optimized geometries.

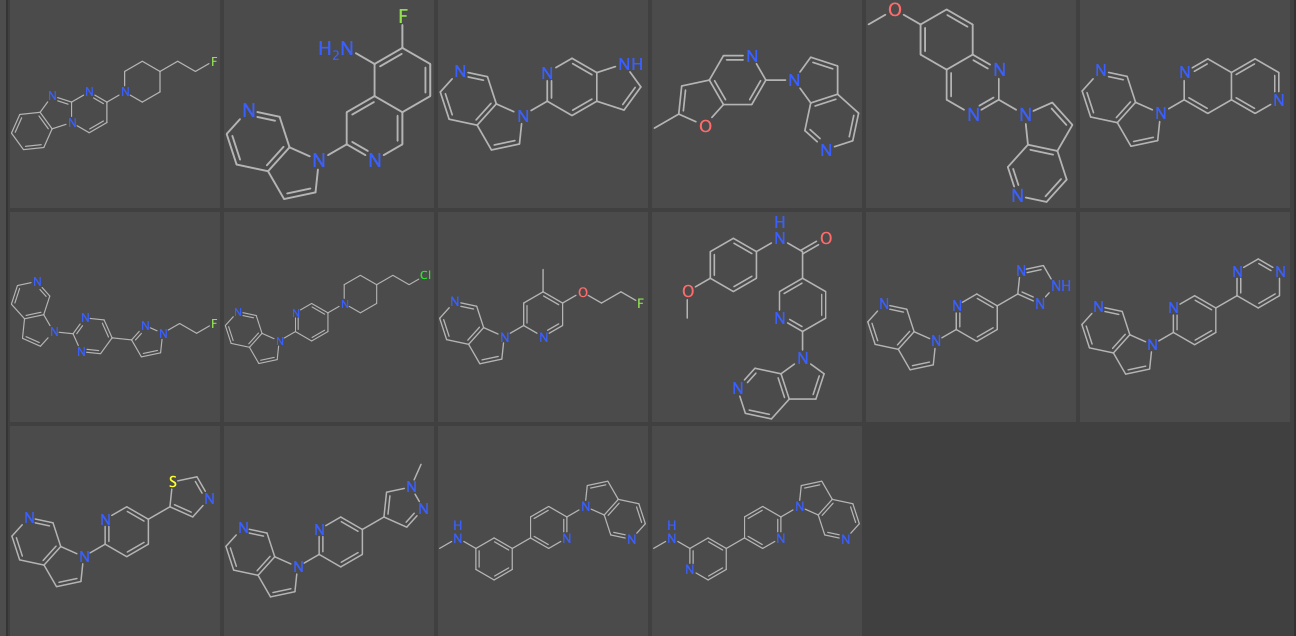
 **Supplemental Table 2.** Known binders against AD PHF tau used for retrospective enrichment. The first listed is GTP-1, and the second is MK-6240. The MK-6240 analogs follow going left to right on each row, named analog_1 through analog_14, in order.

| **Molecule** | **DOCK Score** | **Placement among the 22 Million from ZINC22** |
| --- | --- | --- |
| **GTP-1** | -25.48 | 1454052 |
| **MK-6240** | -26.18 | 1162453 |
| **analog_1** | -22.64 | 3169801 |
| **analog_2** | -25.03 | 1667733 |
| **analog_3** | -29.72 | 308301 |
| **analog_4** | -24.80 | 1785934 |
| **analog_5** | -30.25 | 245840 |
| **analog_6** | -23.55 | 2527669 |
| **analog_7** | -25.10 | 1633332 |
| **analog_8** | -27.63 | 702517 |
| **analog_9** | -28.44 | 517283 |
| **analog_10** | -28.07 | 596019 |
| **analog_11** | -27.60 | 710271 |
| **analog_12** | -28.53 | 499537 |
| **analog_13** | -32.09 | 105140 |
| **analog_14** | -26.90 | 911263 |

**Supplemental Table 3.** Placement of the 16 known binders against AD PHF tau listed in **Supplemental Table 2** among the 22 million ZINC22 molecules docked against the same setup. In total, 12,410,418 molecules out of the 22,037,472 from ZINC22 were able to dock to AD PHF tau in a symmetric way.

**Supplementary Figures**

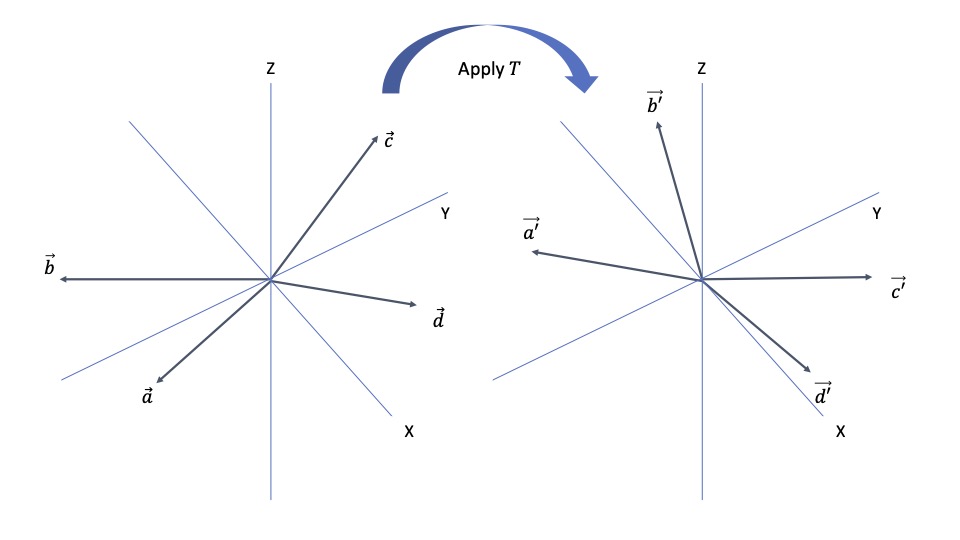

**Supplemental Figure 1.** Determination of an affine transformation matrix from a symmetric system. An affine transformation is a linear transformation (here a rotation) followed by a translation. We can represent translations of vectors in 3 dimensions by matrix multiplication in 4 dimensions by appending a 1 to the vector:

$$\left( \begin{matrix} 1 & 0 & 0 & 0 \\ a & 1 & 0 & 0 \\ b & 0 & 1 & 0 \\ c & 0 & 0 & 1 \end{matrix} \right)\left( \begin{matrix} 1 \\ x \\ y \\ z \end{matrix} \right)=\left( \begin{matrix} 1 \\ a+x \\ b+y \\ c+z \end{matrix} \right).$$

Had we used a rotation matrix $R$ instead of the identity in the lower-right section of the matrix, the action would be rotation by $R$ followed by translation by $\left\{ a,b,c \right\}$. If we have a set of noncoplanar points in 3D $\vec{a}=\{a_{x},a_{y},a_{z}\}$, $\vec{b}=\{b_{x},b_{y},b_{z}\}$, $\vec{c}=\{c_{x},c_{y},c_{z}\}$, and $\vec{d}=\{d_{x},d_{y},d_{z}\}$, and then we apply an affine transformation $T$ to get $\vec{a'}=\{a_{x}^{'},a_{y}^{'},a_{z}^{'}\}$, $\vec{b'}=\{b_{x}^{'},b_{y}^{'},b_{z}^{'}\}$, $\vec{c'}=\{c_{x}^{'},c_{y}^{'},c_{z}^{'}\}$, and $\vec{d'}=\{d_{x}^{'},d_{y}^{'},d_{z}^{'}\}$, respectively, we can relate the initial coordinates to the final ones by

$TA=B$,

where $A=\left( \begin{matrix} 1 & 1 & 1 & 1 \\ a_{x} & b_{x} & c_{x} & d_{x} \\ a_{y} & b_{y} & c_{y} & d_{y} \\ a_{z} & b_{z} & c_{z} & d_{z} \end{matrix} \right)$ and $B=\left( \begin{matrix} 1 & 1 & 1 & 1 \\ a_{x}' & b_{x}' & c_{x}' & d_{x}' \\ a_{y}' & b_{y}' & c_{y}' & d_{y}' \\ a_{z}' & b_{z}' & c_{z}' & d_{z}' \end{matrix} \right)$.

As long as the 4 points are noncoplanar, then we can invert $A$ to find $T$:

$T=BA^{-1}$.

In the case of a symmetric protein, we can find the symmetry operation by selecting 4 noncoplanar points in one copy of the monomer, find their mates in the next, construct the matrices $A$ and $B$, and then compute $BA^{-1}$. Since the protein structure will never be perfectly symmetrical in its deposited coordinates, we did this procedure multiple times for each system to get multiple versions of $T$ and then used the average as the matrix for the whole system.

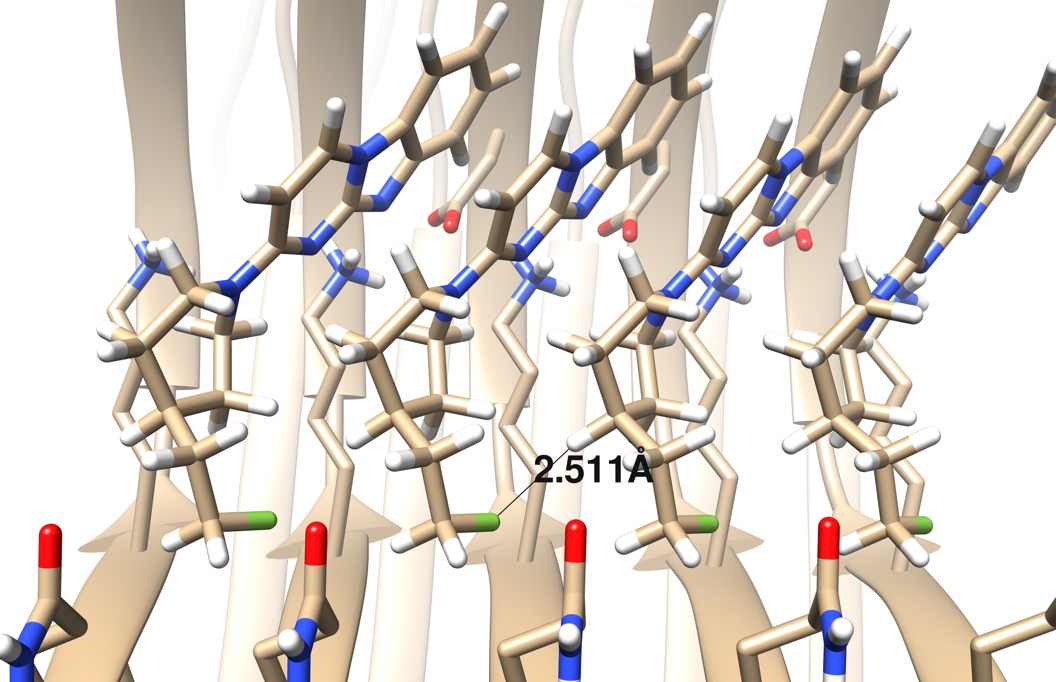

**Supplemental Figure 2.** Determining the cutoff for a clash. In the experimental structure of GTP-1 bound to AD PHF tau shown here, a fluorine of one molecule in the stack is 2.511 Å away from a hydrogen in the next molecule in the stack. The AMBER 4.0 united-atom force field gives the location of zero energy for these two atom types as 2.75 Å, so this distance would be highly unfavorable using this force field. We then take a conservative (and easy to evaluate) definition of a clash as any two atoms on different molecules in the stack being within 2 Å of each other.

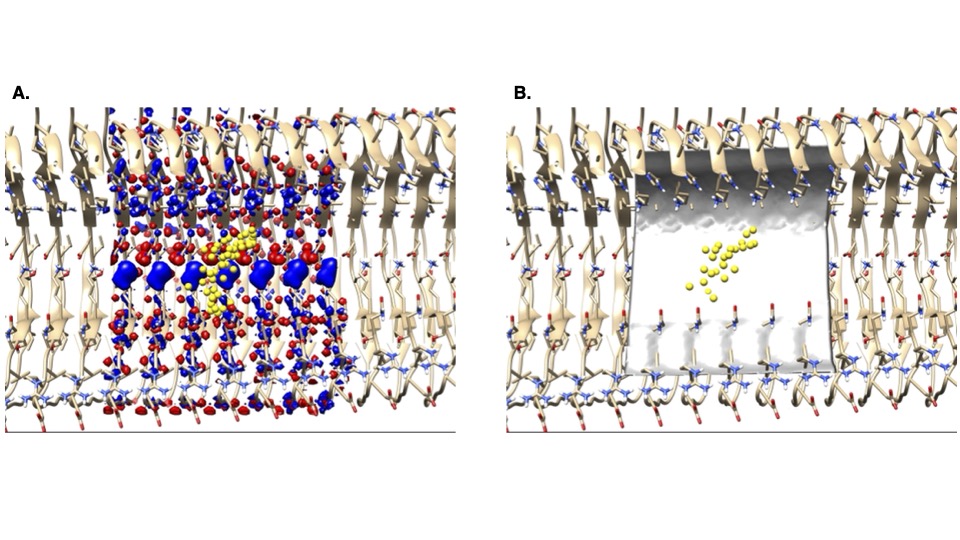

**Supplemental Figure 3.** Using 15 copies of the monomer (GTP-1-bound structure of AD PHF tau, PDB ID: 8FUG) removes edge effects in the (**a.**) electrostatic and (**b.**) desolvation grids in the DOCK score function (yellow for receptor matching spheres to place the ligand, red for an electrostatics level set of -100, blue for an electrostatics level set of 100, and gray for a desolvation level set of 0.768).

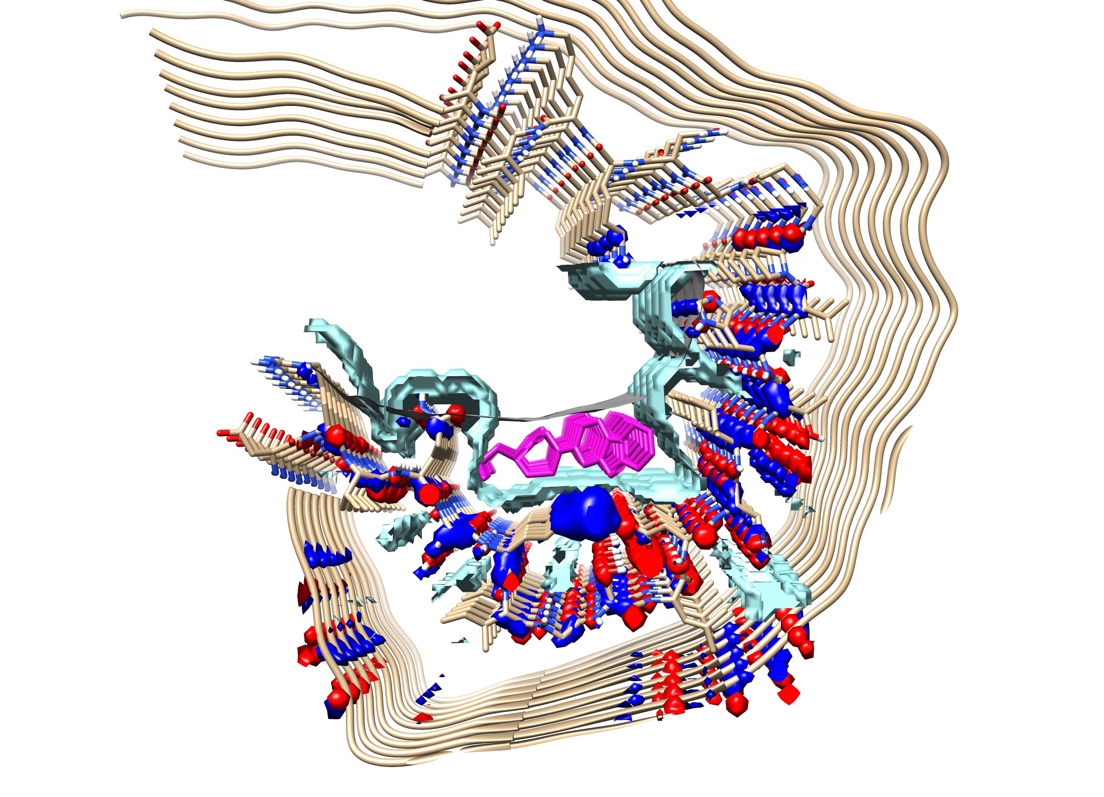

**Supplemental Figure 4.** We account for the changed dielectric environment within a stack of ligands (GTP-1-bound structure of AD PHF tau, PDB ID: 8FUG) by modeling a stack of experimental ligands (magenta) when constructing grids for the electrostatic (red for a level set of -100, blue for a level set of 100) and desolvation (gray for a level set of 0.555, the thin line above the ligand stack), but not van der Waals terms (cyan for a level set of 1.000). This view shows the protein when looking down the fibril.

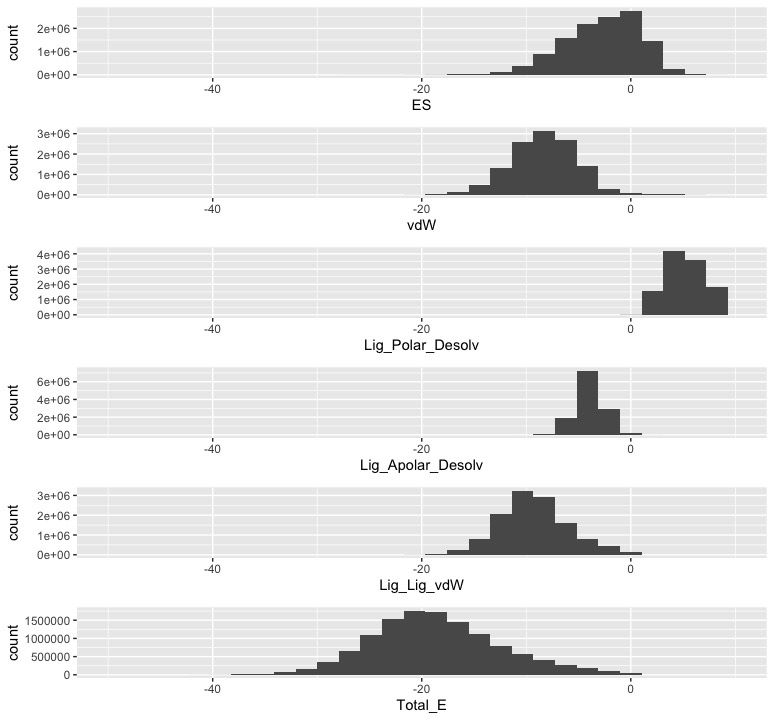

**Supplemental Figure 5.** Effect of changing the low dielectric boundary to account for the ligand stack. The van der Waals component of the DOCK score function is usually the largest in magnitude, and SymDOCK has an extra source of van der Waals terms coming from the ligand-ligand interactions. These two energy sources tend to dominate the total score to the detriment of the electrostatic and ligand desolvation terms, weakening the effects of specific interactions with the protein. To overcome this imbalance, we increase the region of low dielectric when calculating the electrostatic and ligand desolvation grids to include the region where a ligand stack would go. Decreasing the dielectric in these terms increases their magnitude, putting them more in line with the (2) van der Waals terms. To test this balancing, we broke down the score components of docking the 22.0 million molecules from ZINC22 as described in the main text. We show these as counts in each energy bin and see that the electrostatic and ligand desolvation values are closer to the van der Waals terms.

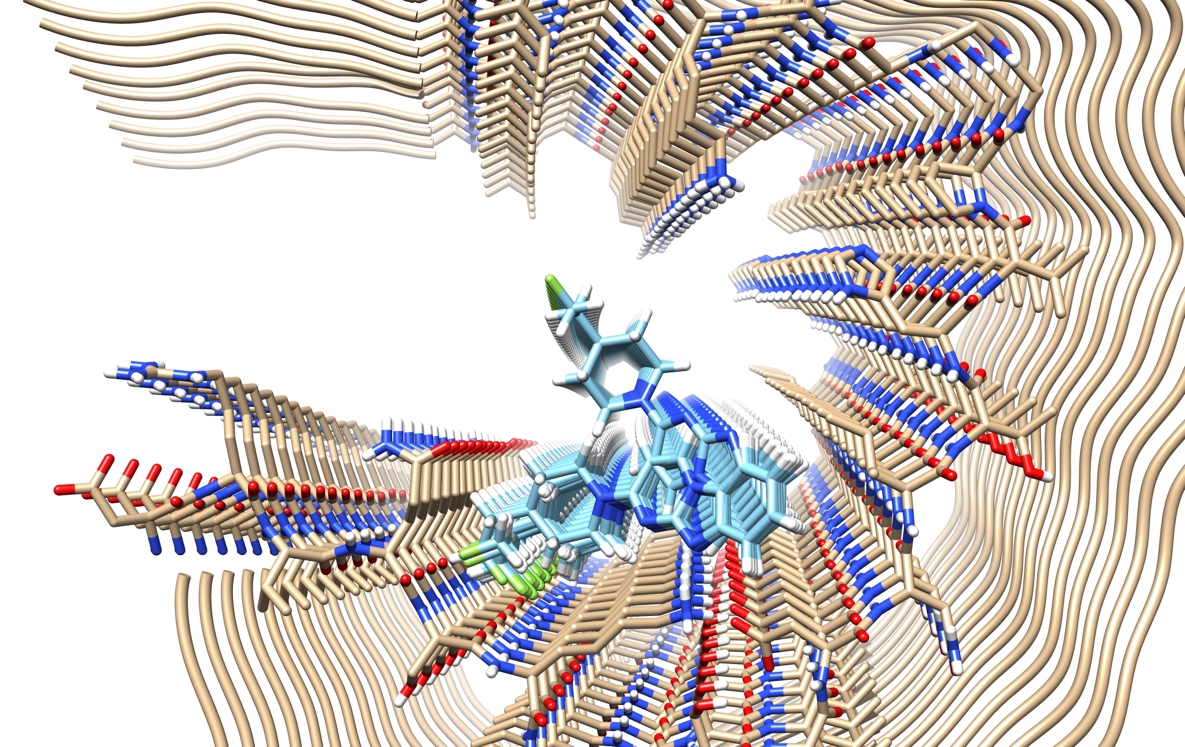

**Supplemental Figure 6.** Top 20 (by DOCK score) alternate docked poses of GTP-1 (blue) against AD PHF tau (beige).

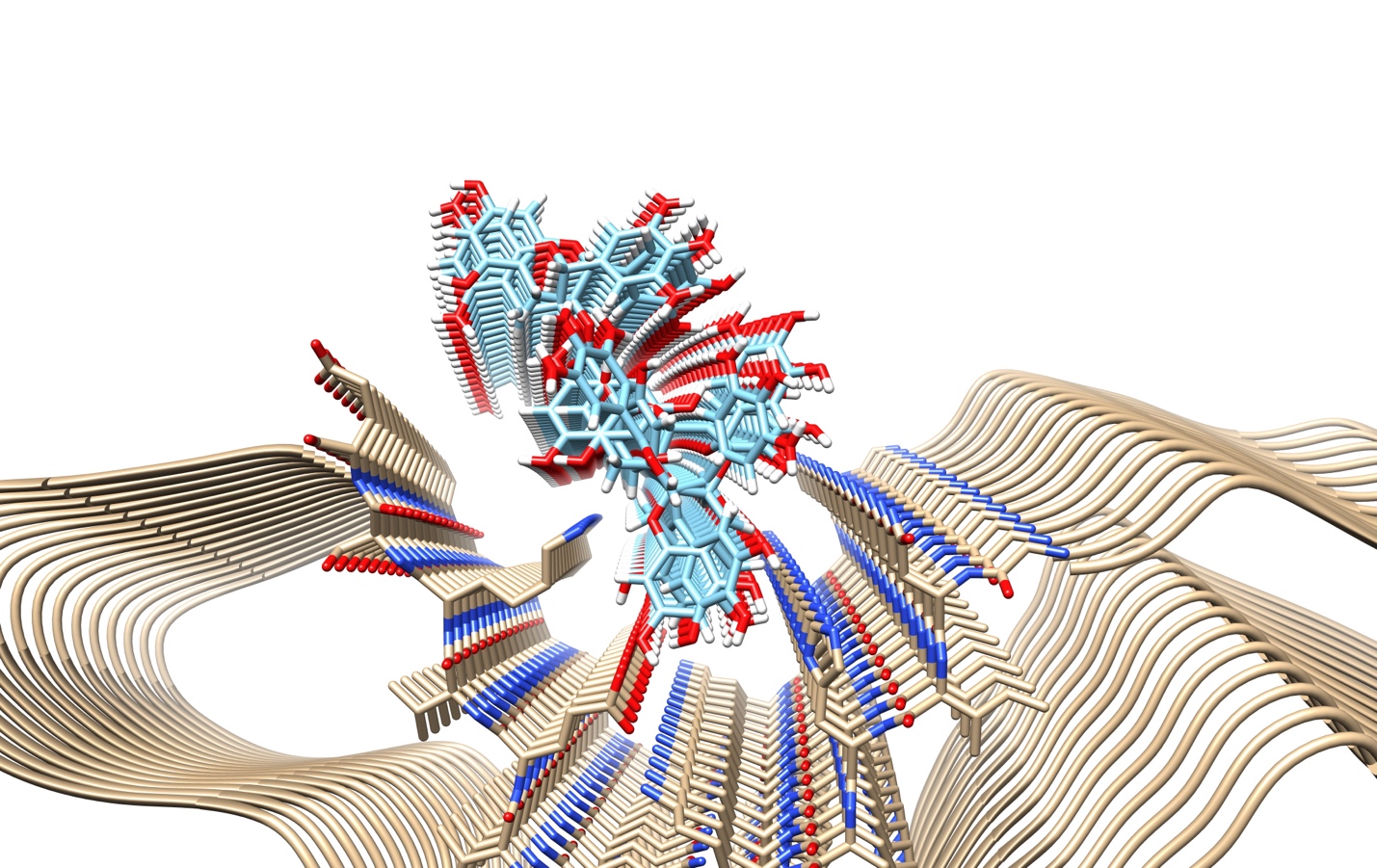

**Supplemental Figure 7.** Top 50 (by DOCK score) alternate docked poses of EGCG (blue) against AD PHF tau (beige).

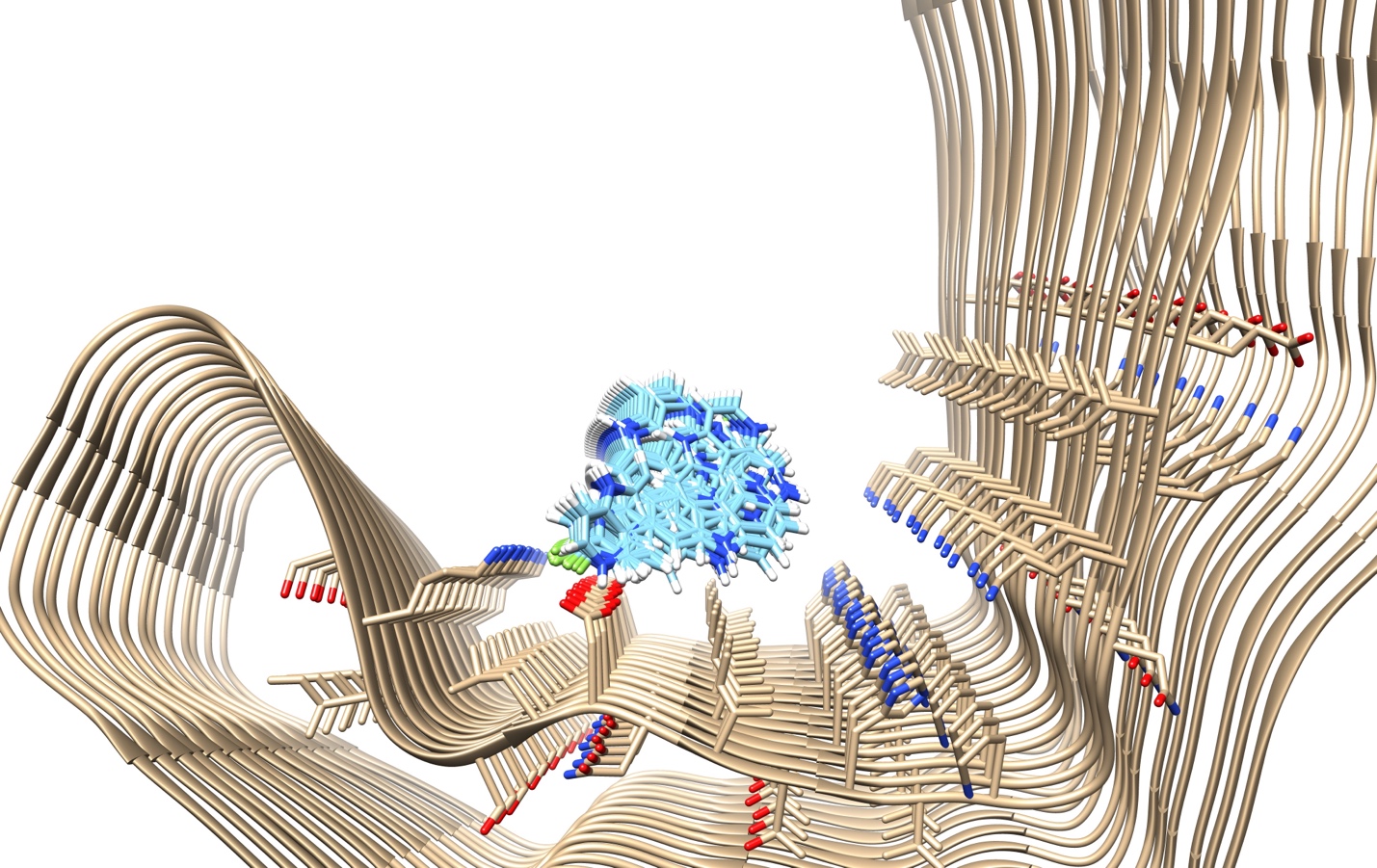

**Supplemental Figure 8.** Top 50 (by DOCK score) alternate docked poses of flortaucipir (blue) against CTE Type I tau (beige).

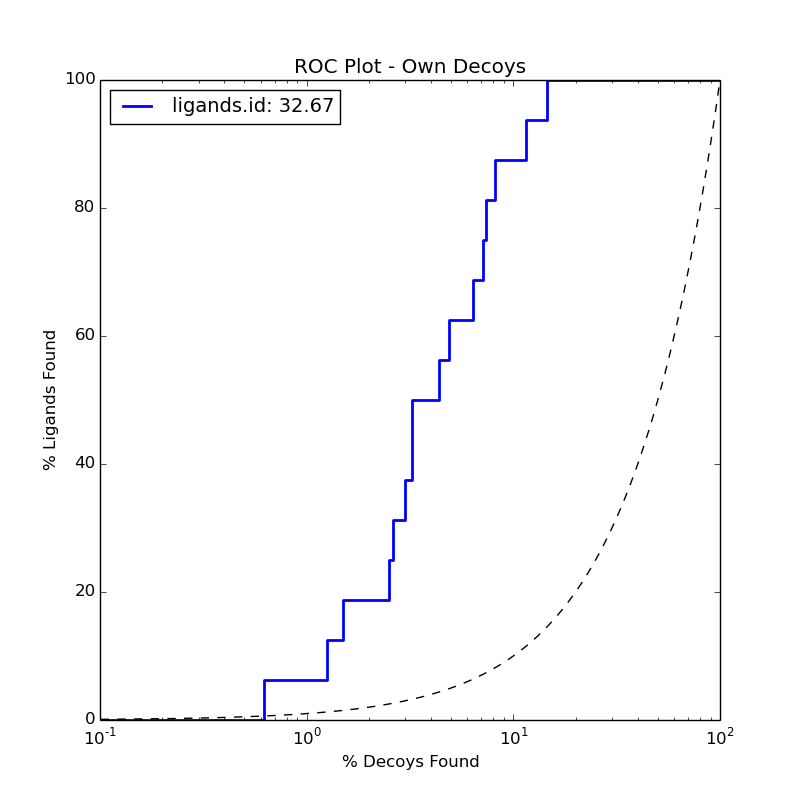

**Supplemental Figure 9.** Enrichment of the 16 known binders above of AD PHF tau against 800 property-matched decoys using SymDOCK, giving an adjusted logAUC enrichment of 32.67 and an enrichment factor at 1% (EF_1%_) of 6.375.

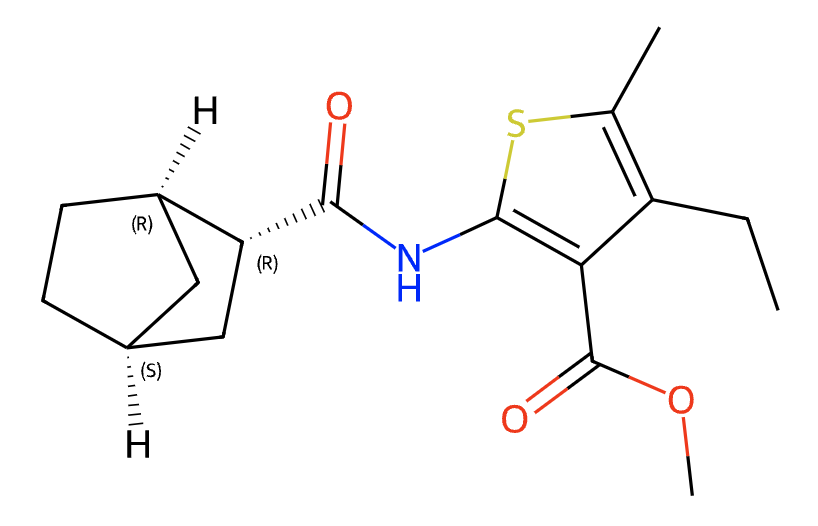

**Supplemental Figure 10.** Structure of the molecule ZINC000000035542, a decoy used for the retrospective enrichment study of AD PHF tau. This molecule could dock to the fibril using the default DOCK3.8 procedure but not using SymDOCK.

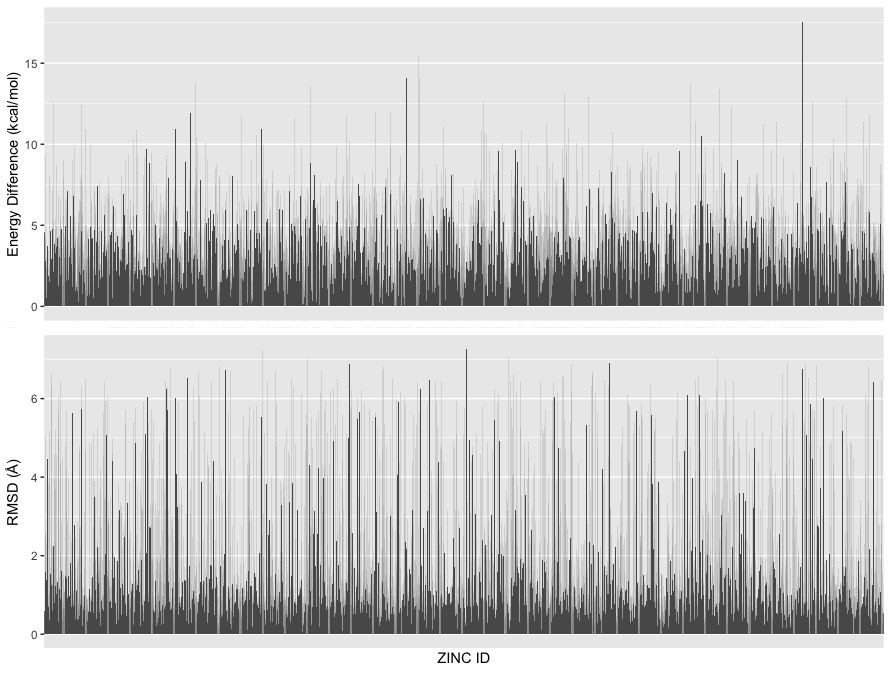

**Supplemental Figure 11.** Comparison of the SymDOCK-generated poses to ANI-optimized geometries among the top 5000 (by DOCK score) molecules from docking 22.0 million from ZINC22. Top: difference in energy (energy for SymDOCK pose passed through ANI minus ANI energy for optimized pose). Bottom: root-mean-square distance between the SymDOCK pose and ANI-optimized pose.
